# MOSPD3 mediates the ER recruitment of ATG2A to promote autophagy

**DOI:** 10.64898/2026.01.04.697522

**Authors:** Yuanjiao Du, Tiantian Zhou, Yazhou Liu, Chunyu Song, Wei-Ke Ji

## Abstract

ATG2A transfers glycerophospholipids from the endoplasmic reticulum (ER) to support expansion of the isolation membrane at ER membrane contact sites. While a role for ATG18/WIPI4 in targeting ATG2A to the IM is well established, how ATG2A is recruited to the ER remains unclear. In this study, we found that MOSPD3, an atypical VAP family protein, acts an adaptor that recruits ATG2A to the ER. MOSPD3 colocalizes with ATG2A and is specifically enriched at ER sites juxtaposed with the IM during autophagosome formation. Recruitment is mediated by direct interactions between the FFNT (two phenylalanines in a neutral tract) motif in the N-terminal region (NT) of ATG2A and the major sperm protein (MSP) domain of MOSPD3. Co-expression of MOSPD3 with ATG2A-NT, but not a MOSPD3-binding-defective mutant ATG2A-NT-T362A, markedly rescued the autophagic defect caused by ATG2A/B double knockout (DKO). MOSPD3 depletion abolishes ATG2A recruitment to the ER and impedes autophagic flux. Together, this study demonstrates that MOSPD3 functions as an ER adaptor for ATG2A in autophagy.

## Introduction

Macroautophagy (hereafter referred to as autophagy) is an intracellular quality control process that is highly conserved in eukaryotes. This process involves on the de novo biogenesis of an isolation membrane (IM), a cup-shaped membrane sac, and its further expansion^1^. The IM eventually seals to form a double-membrane autophagosome that fuses with a lysosome to degrade engulfed cytoplasmic toxins.

Although multiple organelles have been proposed as potential membrane sources for autophagosome biogenesis^2^, it is generally accepted that the endoplasmic reticulum (ER) plays an essential role in autophagosome biogenesis^3–6^. In terms of the evidence supporting this, the ER is known to form extensive contacts with the IM during development into an autophagosome, and the IM is embedded between closely spaced ER cisternae and connected to the associated ER via membrane extensions, which dissociate from the ER after it matures into an autophagosome^7, 8^.

ATG2A/B are core ATG proteins that act in the early stage of autophagosome maturation^9, 10^. ATG2A/B are similar in structure to the VPS13 proteins by also exhibiting an extended hydrophobic groove along their entire length and belong to the bridge-like transfer protein (BLTP) family^11, 12^. Previous studies have shown that ATG2A is specifically localized at ER–IM contacts, physically bridging the gap between the edge of the expanding IM and the ER. At these sites, ATG2 proteins efficiently transfer glycerophospholipids from the ER to serve as building blocks for autophagosome formation^13–15^, in cooperation with the lipid scramblases TMEM41B/VMP1 and ATG9^16, 17^. The targeting of ATG2 to IM relies on its interactions with ATG18/WIPI4 and ATG9, and between ATG18/WIPI4 and phosphatidylinositol 3-phosphate (PI3P)^17–22^. Although the tethering of ATG2 to ER membranes is a key step in commitment to the autophagosome formation pathway^23^, it remains unclear how ATG2A/B are recruited to the ER.

Given the established role of the FFAT motif (diphenylalanine [FF] in an acidic tract) in the recruitment of soluble lipid transfer proteins such as VPS13 proteins^24, 25^, we hypothesized that ATG2 proteins may associate with the ER via FFAT or FFAT-like motifs that bind directly to vesicle-associated membrane protein-associated proteins (VAPs) on the ER^26, 27^. In this study, we tested this hypothesis by performing targeted screening to investigate whether VAP and VAP-related proteins, including VAPA, VAPB, MOSPD2, MOSPD1, and MOSPD3, are capable of recruiting ATG2 proteins to the ER.

## Results

### MOSPD3 interacts with ATG2A and is specifically required for its ER association

In the targeted screening, we investigated colocalization between GFP-ATG2A and each VAP protein. Notably, ectopically overexpressed GFP-ATG2A was mainly found on lipid droplets (LDs) labeled by BODIPY-C12, but not on IMs (Fig. S1A), as previously reported^14, 28^. Importantly, we found that MOSPD3 (Fig. S1B), but not the other VAP-related proteins (Fig. S1C–F, S1G), strongly colocalized with GFP-ATG2A, possibly at ER–LD junctions, suggesting that MOSPD3 is an ER adaptor of ATG2A.

To avoid the artificial effect of ectopic GFP-ATG2A, we investigated the spatial relationship between endogenous ATG2A and MOSPD3 in a U2OS cell line with CRISPR–Cas9-mediated knock-in of GFP in the C-terminal of ATG2A (ATG2A-GFP-KI)^29^. First, co-immunoprecipitation (coIP) showed that MOSPD3 interacted with ATG2A at endogenous levels in this line (Fig. 1A).

**Fig. 1.**
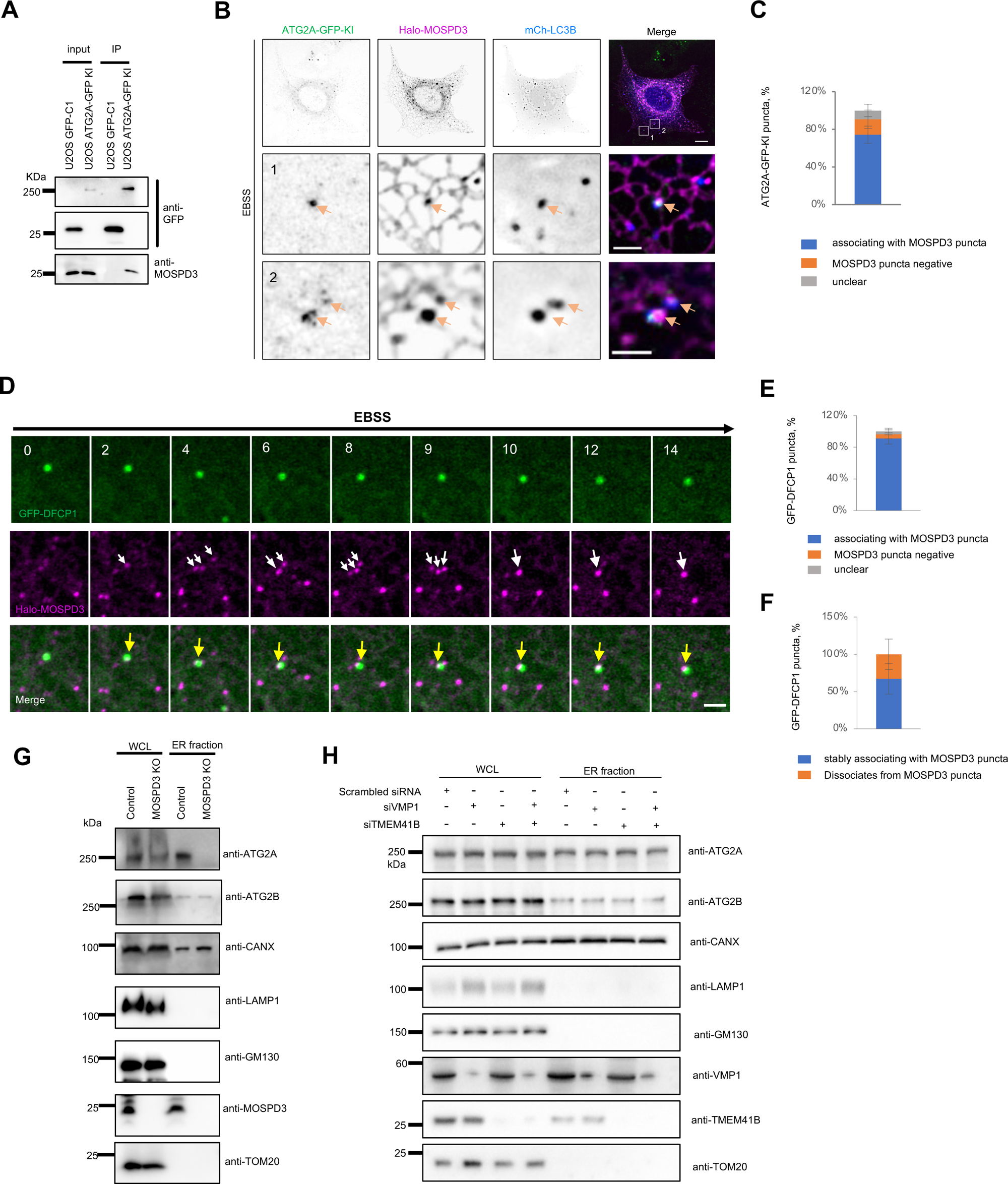
MOSPD3 interacts with ATG2A and is specifically required for ER association of ATG2A. **a.** Representative GFP-Trap assays demonstrating an interaction between endogenous ATG2A-GFP-KI and endogenous MOSPD3 from at least 3 independent experiments. **b.** Representative images of a live U2OS ATG2A-GFP KI cell transiently transfected with Halo-MOSPD3 (magenta) and mCh-LC3B (blue) with insets under EBSS stimulation. **c.** The percentage of endogenous ATG2A puncta that colocalized with MOSPD3 puncta (832 ATG2A-GFP puncta from 15 cells) from at least 3 independent experiments. Mean ± SD. **d.** Time-lapse montages of a COS7 cell transiently transfected with GFP-DFCP (green) and Halo-MOSPD3 (magenta) with white arrows denoting dynamic MOSPD3 puncta and yellow arrows denoting colocalization of GFP-DFCP1 puncta and Halo-MOSPD3 puncta from 3 independent experiments. **e.** The percentage of DFCP1 puncta relative to MOSPD3 puncta from 12 cells. Mean ± SD. **f.** The percentage of DFCP1 puncta stably associating with MOSPD3 puncta from 12 cells. GFP-DFCP1 puncta associating with Halo-MOSPD3 puncta over the imaging time window (∼15 min) were defined to be stable association. Mean ± SD. **g.** Membrane fractionation showing the ER association of ATG2A and ATG2B in WT and MOSPD3 KO HeLa cells in 3 independent experiments. Western blots were performed with antibodies against ATG2A, ATG2B, CANX (ER marker), Lamp1 (late endosome/PM marker), GM130 (Golgi marker) and TOM20 (mitochondrial marker). **h.** Membrane fractionation showing the ER association of ATG2A and ATG2B in HeLa cells treated with scrambled, VMP1, TMEM41B or VMP1+TMEM41B siRNAs in 3 independent experiments. Western blots were performed with antibodies against ATG2A, ATG2B, VMP1, TMEM41B, CANX (ER marker), Lamp1 (late endosome/PM marker), GM130 (Golgi marker) and TOM20 (mitochondrial marker). Scale bar, 10 μm in the whole cell image, 2 μm in the insets in (b, d).

Second, we found that MOSPD3 formed puncta on the ER, likely representing its enrichment in ER subdomains (Fig. 1B). Notably, a significant portion of ATG2A-GFP-KI puncta (>70%) were well colocalized with these MOSPD3 puncta and appeared to be present at the contacts between the ER and IMs marked by LC3B upon autophagic induction (Fig. 1B, C; Fig. S2A, B).

Next, we tracked the spatial and temporal relationship between MOSPD3 and IMs labeled with DFCP1, which forms puncta immediately after autophagy activation^30, 31^. Time-lapse video analysis showed that MOSPD3 was able to form puncta in response to starvation, and some small MOSPD3 puncta could fuse at sites where DFCP1 puncta was present, possibly representing autophagosome formation sites on the ER (Fig. 1D). Live imaging revealed that most GFP-DFCP1 puncta colocalized with MOSPD3 puncta (Fig. 1E), and a significant portion of the association was stable over time upon autophagic induction (Fig. 1F).

To investigate whether MOSPD3 is required for the recruitment of ATG2A to the ER, we performed cell fractionation to isolate the ER fraction from either control or CRISPR–Cas9-mediated MOSPD3 knockout (KO) HeLa cells (Fig. S2C), which was confirmed by genotyping and western blotting (Fig. S2D, E). We confirmed the purity of the ER fractions and found that MOSPD3 was specifically required for the ER association of ATG2A, as the ER-associated pool of ATG2A was almost completely lost in MOSPD3-KO cells, whereas that of ATG2B was unaffected (Fig. 1G). These results indicate that MOSPD3 plays an essential role in the recruitment of ATG2A to the ER.

Lipid scramblases, such as ER-resident VMP1 and TMEM41B, and IM-resident ATG9, are thought to mediate lipid equilibration between two leaflets of ER and IM membranes and participate in the organization of ER–IM contacts during autophagosome maturation^16, 17, 32^. Given the close functional relationship between ATG2 and these scramblases, we investigated whether the ER-resident VMP1 and TMEM41B could mediate the recruitment of ATG2 to the ER. In contrast to the clear colocalization between ATG2A and MOSPD3 (Fig. S3A), neither TMEM41B nor VMP1 showed significant colocalization with ATG2A under normal (Fig. S3B, C) or starvation conditions (Fig. S3D). In addition, we found that an N-terminal (NT) region of ATG2A (ATG2A-NT), which was responsible for interacting with MOSPD3 (Fig. S3E, top panel; also see the next result section), was not recruited to the ER even after overexpression of these two lipid scramblases (Fig. S3E, F).

Consistent with imaging results, small interfering RNA (siRNA)-mediated depletion of TMEM41B or VMP1, individually or combinatorial, did not reduce the levels of ATG2A or ATG2B in the ER membrane fraction (Fig. 1H). Therefore, these results indicate that VMP1 and TMEM41B are not ER adaptors of ATG2, even though these two proteins are functional partners of ATG2.

### An FFNT motif in ATG2A is required for its recruitment by MOSPD3

We next attempted to clarify how ATG2A is recruited by MOSPD3 via molecular dissection. We found that an ATG2A mutant lacking a C-terminal region reported to mediate association with phagophores or LDs^14, 28^ was cytosolic (Fig. 2A; inset 1). In contrast, this truncation mutant could be recruited to the ER in a cell transfected with Halo-MOSPD3 in the same field of view (Fig. 2A; inset 2).

**Fig. 2.**
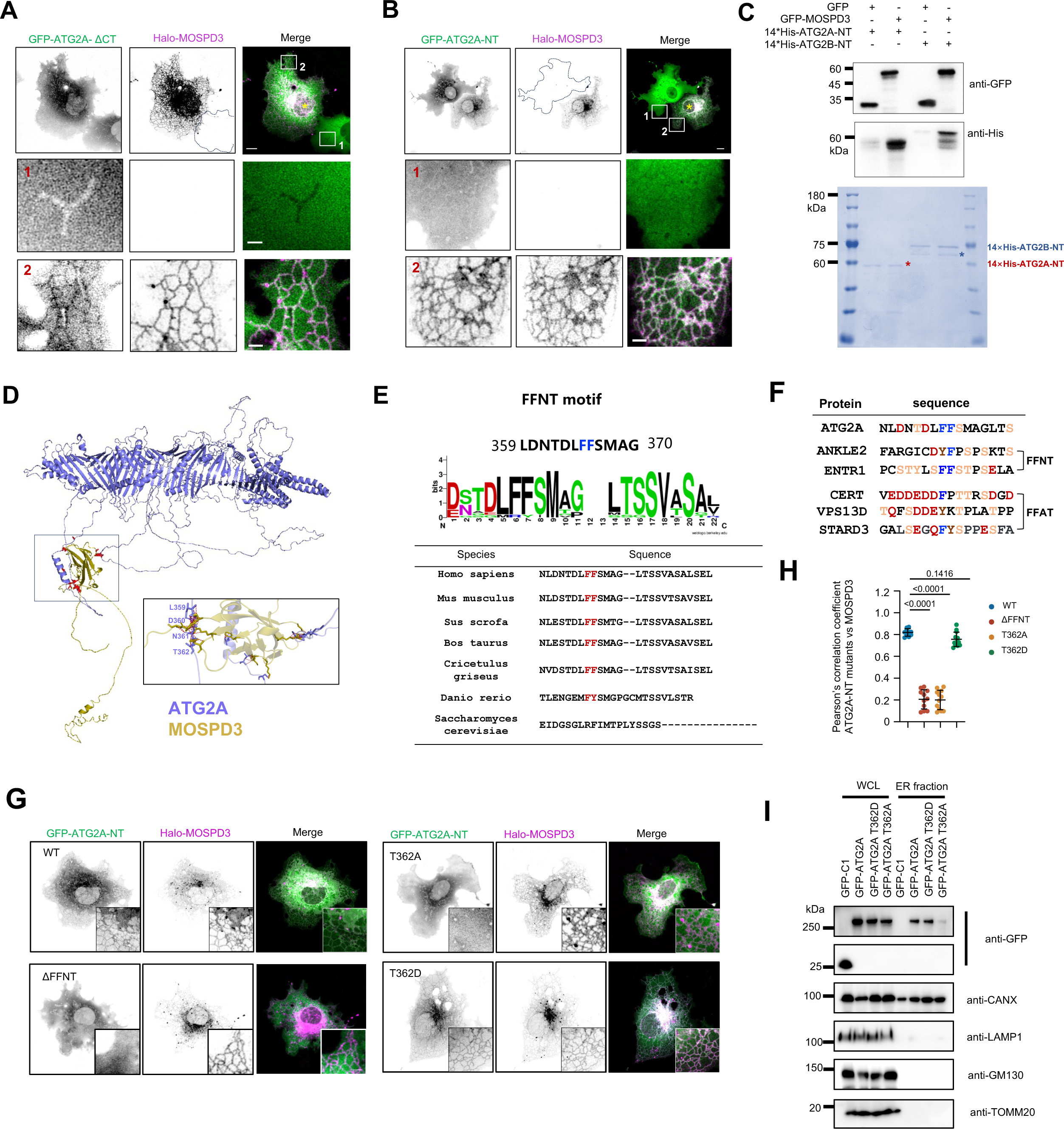
An FFNT motif in ATG2A-NT is required for ER recruitment of ATG2A by MOSPD3. **a.** Representative images of a COS7 cell expressing GFP-ATG2A-ΔCT (green) and Halo-MOSPD3 (magenta) with two insets from at least 3 independent experiments. Yellow asterisks mark cells expressing both GFP-ATG2A-ΔCT and Halo-MOSPD3. **b.** Representative images of a COS7cell expressing GFP-ATG2A-NT (green) and Halo-MOSPD3 (magenta) with two insets from at least 3 independent experiments. Yellow asterisks mark cells transfected with both GFP-ATG2A-ΔCT and Halo-MOSPD3. **c.** In vitro pulldown assays using purified GFP-MOSPD3 and purified 14×His-ATG2A-NT or 14×His-ATG2B-NT from 3 independent experiments. **d.** AlphaFold Multimer prediction of the binding sites between ATG2A and MOSPD3. **e.** Conservation of the FFNT motif in ATG2A across species. **f.** The conserved FFNT motif within ATG2A is compared with homologous regions from the indicated human proteins (ANKLE2, CERT, VPS13D, and STARD3). Conserved residues are highlighted. **g.** Representative images of COS7 cells expressing GFP-ATG2A-NT (green; WT, T362A, T362D and the ΔFFN mutant) and Halo-MOSPD3 (magenta) with insets from at least 3 independent experiments. **h.** Pearson’s correlation coefficient of MOSPD3 vs. WT ATG2A-NT (10 cells), ATG2A-NT-ΔFFNT (11 cells), ATG2A-T362A (12 cells), or ATG2A-T362D (12 cells) in three independent experiments. Ordinary one-way ANOVA with Tukey’s multiple comparisons test. Mean ± SD. **i.** Membrane fractionation showing the ER association of WT GFP-ATG2A-NT, GFP-ATG2A-NT-T362A and GFP-ATG2A-NT-T362D from 3 independent experiments. Western blots were performed with antibodies against GFP, CANX (ER marker), Lamp1 (late endosome/PM marker), GM130 (Golgi marker) and TOM20 (mitochondrial marker). Scale bar, 10 μm in the whole cell images and 2 μm in the insets in (a, b, & g).

We also expressed an ATG2A truncation mutant containing part of the N-terminal region (residues 1–380; ATG2A-NT) with Halo-MOSPD3. Notably, ATG2A-NT was cytosolic (Fig. 2B; inset 1) but could be strongly recruited to the ER in cells expressing Halo-MOSPD3 in the same field of view (Fig. 2B; inset 2). These results suggest that the ATG2A-NT is responsible for interaction with MOSPD3. Indeed, in vitro pulldown assays indicated direct interaction between purified ATG2A-NT and GFP-MOSPD3, but not with GFP tag alone (Fig. 2C). Notably, MOSPD3 also bound to an NT region of ATG2B (ATG2B-NT, residues 1-450) but to a much lesser extent than the ATG2A-NT.

As a member of the VAP-related proteins, MOSPD3 likely recognizes an FFNT motif in ATG2A, and the binding suggests the presence of an FFAT or FFAT-like motif in ATG2A-NT. To identify the position of the FFAT or FFAT-like motif in ATG2A-NT, we attempted to predict protein–protein interactions using AlphaFold Multimer^33^. The predicted models consistently indicated that a largely disordered region in ATG2A-NT outside the hydrophobic groove mediates the interaction with the MSP domain of MOSPD3 (Fig. 2D). Notably, a short sequence (residues 359 to 370) in this region showed similarity to the canonical FFAT motif but with a smaller number of acidic residues around two consecutive FF residues (Fig. 2E), consistent with the feature of the FFNT motif (diphenylalanine [FF] in a neutral tract) first proposed by Cabukusta et al.^34^. The FFNT motif was highly conserved at the amino acid level among vertebrates but not in yeast (Fig. 2E, F).

In contrast to wild-type (WT) ATG2A-NT, a mutant lacking the putative FFNT motif failed to target to the ER, even when Halo-MOSPD3 was expressed (Fig. 2G, left panel). Remarkably, a point mutant in the FFNT motif of ATG2A-NT (ATG2A-NT-T362A) could not be recruited to the ER by Halo-MOSPD3, whereas replacement of T362 with a polar aspartate residue (T362D) still could (Fig. 2G, right panel; 2H). These results suggest that the FFNT motif is required for ER recruitment of ATG2A by MOSPD3 and T362 is essential for the integrity and function of this FFNT motif.

We next performed cell fractionation to confirm the importance of the FFNT motif in the ER recruitment of ATG2A. Consistent with the imaging data, the ER pool of GFP-ATG2A-T362A was evidently reduced compared with that of the WT or GFP-ATG2A-T362D mutant (Fig. 2I).

### The MSP domain of MOSPD3 mediates the recruitment of ATG2A to the ER

MOSPD3 contains an MSP domain in the NT and a proline-rich region followed by two consecutive transmembrane (TM) domains (Fig. 3A). A MOSPD3 mutant lacking the two TMs was diffuse in the cytosol but could be recruited to GFP-ATG2A (Fig. 3B), suggesting a reversible recruitment between MOSPD3 and ATG2A.

**Fig. 3.**
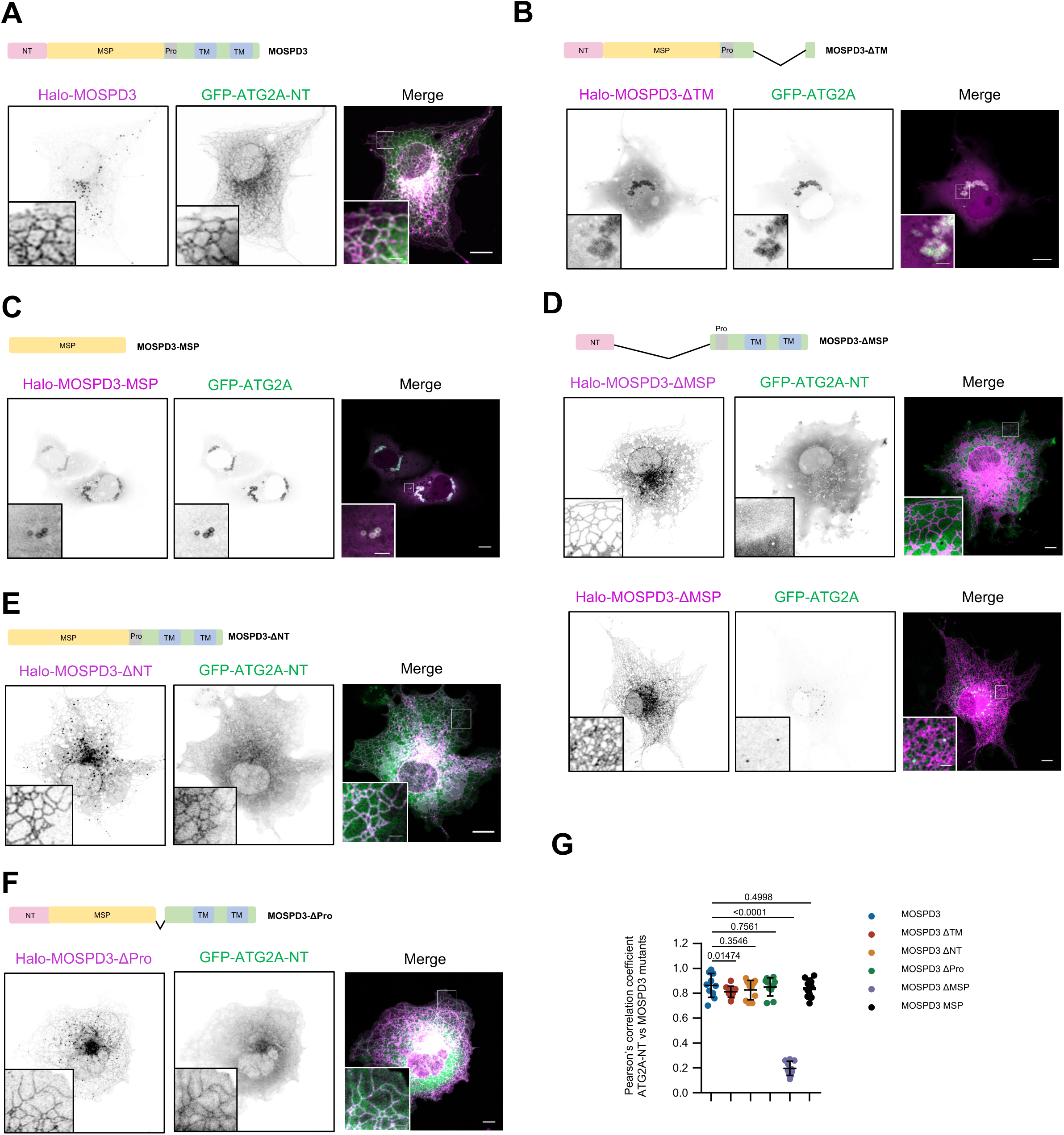
The MSP domain in MOSPD3 mediates ATG2A recruitment to the ER. **a.** Top: domain organization of MOSPD3. Bottom: representative images of a COS7 cell expressing GFP-ATG2A-NT (green) and Halo-MOSPD3 (magenta) with insets from at least 3 independent experiments. **b.** Representative images of a COS7 cell expressing GFP-ATG2A (green) and Halo-MOSPD3-ΔTM (magenta) with insets from at least 3 independent experiments. **c.** Representative images of a COS7 cell expressing GFP-ATG2A (green) and Halo-MOSPD3-MSP (magenta) with insets from at least 3 independent experiments. **d.** Representative images of a COS7 cell expressing either GFP-ATG2A-NT (green; top panel) or GFP-ATG2A (green; bottom panel) along with Halo-MOSPD3-ΔMSP (magenta) with insets from at least 3 independent experiments. **e.** Representative images of a COS7 cell expressing GFP-ATG2A-NT (green) and Halo-MOSPD3-ΔNT (magenta) with insets from at least 3 independent experiments. **f.** Representative images of a COS7 cell expressing GFP-ATG2A-NT (green) and Halo-MOSPD3-ΔPro (magenta) with insets from at least 3 independent experiments. **g.** In cells as in (**a-f**), Pearson’s correlation coefficient of ATG2A vs different MOSPD3 mutants including WT Halo-MOSPD3, Halo-MOSPD3-ΔNT, Halo-MOSPD3-Δ Pro, Halo-MOSPD3-ΔMSP, Halo-MOSPD3-ΔTM, Halo-MOSPD3-MSP. 10 cells for each group were analysed from three independent experiments. Ordinary one-way ANOVA with Tukey’s multiple comparisons test. Mean ± SD. Scale bar, 10 μm in the whole cell images and 2 μm in the insets in (a-f).

Consistent with the MSP domain playing a key role in the recruitment of FFAT/FFNT-containing proteins, a MOSPD3 truncation mutant containing only the MSP domain was strongly recruited to GFP-ATG2A (Fig.3C), and a mutant lacking MSP (MOSPD3-ΔMSP) abolished the colocalization with GFP-ATG2A-NT (Fig. 3D, top panel) or GFP-ATG2A (Fig. 3D, bottom panel).

In addition, MOSPD3 truncation mutants lacking either an N-terminal region upstream of the MSP domain (MOSPD3-ΔNT; Fig. 3E) or the proline-rich region (MOSPD3-ΔPro; Fig. 3F, G) did not obviously affect the recruitment of ATG2A-NT to the ER. Taking these findings together, MOSPD3 recruits ATG2A via interaction between the MSP domain of MOSPD3 and an FFNT motif of ATG2A.

### ATG2B is recruited to the ER via multiple VAPs including MOSPD1 or MOSPD3

We next investigated the ER recruitment mechanism of ATG2B. Unlike GFP-ATG2A, ectopically expressed GFP-ATG2B exhibited diffuse cytosolic localization (Fig. 4A). Strikingly, co-expression with either Halo-MOSPD1 or Halo-MOSPD3 but not with other VAP proteins, efficiently recruited GFP-ATG2B to the ER (Fig. 4A, B).

**Fig. 4.**
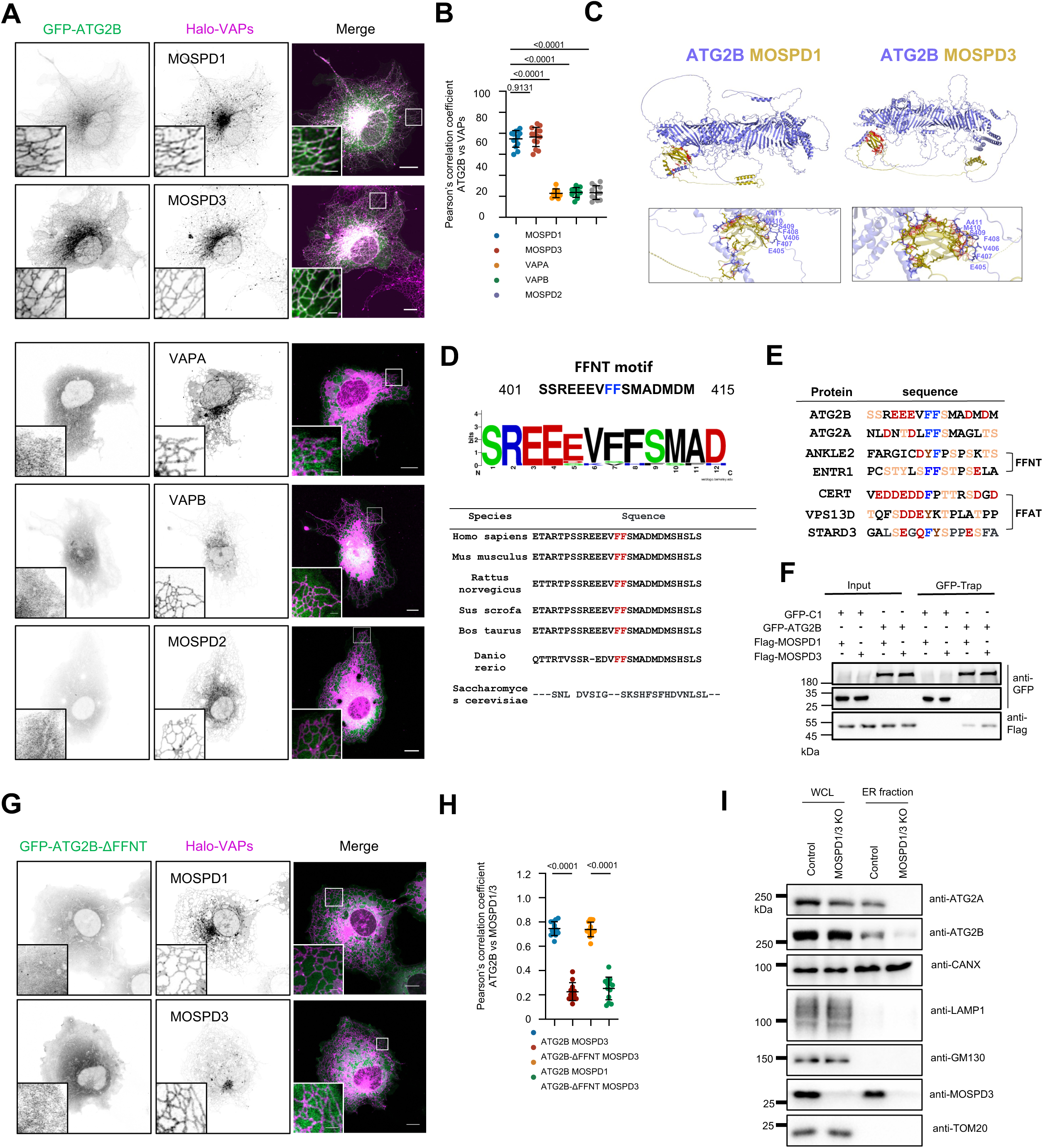
ATG2B is recruited to the ER via multiple VAPs. **a.** Representative images of COS7 cells transiently transfected with GFP-ATG2B (green) along with MOSPD1, MOSPD3, VAPA, VAPB or MOSPD2 (magenta) with an inset on the bottom from at least 3 independent experiments. **b.** In cells as in (**a**), Pearson’s correlation coefficient of ATG2B vs different VAP-related proteins including Halo-MOSPD1 (13 cells), Halo-MOSPD3 (14 cells), Halo-MOSPD2 (12 cells), Halo-VAPA (10 cells), or Halo-VAPB (11 cells) in three independent experiments. Ordinary one-way ANOVA with Tukey’s multiple comparisons test. Mean ± SD. **c.** AlphaFold Multimer prediction of the binding sites between ATG2B and MOSPD1 (left panel) or MOSPD3 (right panel). **d.** Conservation of the FFNT motif in ATG2B across species. **e.** The conserved FFNT motif within ATG2B is compared with homologous regions from the indicated human proteins (ATG2A, ANKLE2, ENTR1, CERT, VPS13D, and STARD3). Conserved residues are highlighted. **f.** GFP-Trap assays demonstrate interactions between ATG2B and MOSPD1/3 in HEK293 cells from three independent assays. **g.** Representative images of COS7 cells expressing GFP-ATG2B-ΔFFNT (green) along with either Halo-MOSPD1 (magenta; top panel) or Halo-MOSPD3 (magenta; bottom panel) with insets from at least 3 independent experiments. **h.** In cells as in (**g**), Pearson’s correlation coefficient of ATG2B-ΔFFNT vs Halo-MOSPD1 (12 cells), or Halo-MOSPD3 (13 cells) in three independent experiments. Unpaired student t test. Mean ± SD. **i.** Membrane fractionation showing the ER association of ATG2A and ATG2B in MOSPD1/3 double KO HeLa cells from 3 independent experiments. Western blots were performed with antibodies against ATG2A, ATG2B, MOSPD3, CANX (ER marker), Lamp1 (late endosome/PM marker), GM130 (Golgi marker) and TOM20 (mitochondrial marker). Scale bar, 10 μm in the whole cell images and 2 μm in the insets in (a & g).

AlphaFold Multimer prediction suggested that ATG2B interacted with MOSPD1 and MOSPD3 via a short linear sequence outside its hydrophobic groove (Fig. 4C). This segment, which is structurally similar to the FFNT motif in ATG2A, is conserved among vertebrates (Fig. 4D, E). Consistent with these predictions, co-IP assays confirmed specific interactions between ATG2B and MOSPD1 or MOSPD3 (Fig. 4F).

These predictions were further supported by the findings of colocalization analyses, in which an ATG2B mutant lacking the predicted FFNT motif (GFP-ATG2B-ΔFFNT) could not be recruited to the ER by either Halo-MOSPD1 or Halo-MOSPD3 (Fig. 4G, H). This confirms that the FFNT motif is essential for MOSPD1/3-mediated ER targeting of ATG2B.

Given that MOSPD3 depletion alone did not impair ATG2B localization to the ER (Fig. 1G), we hypothesized that MOSPD1 and MOSPD3 function redundantly in this process. To test this, we generated a MOSPD1/3 double-knockout (DKO) HeLa cell line (Fig. S2F–I). Cell fractionation revealed a substantial reduction in the ER-associated pool of GFP-ATG2B in the DKO cells compared to wild-type controls (Fig. 4I). Notably, a residual ER-associated pool persisted, suggesting that additional mechanisms may contribute to ATG2B recruitment. This partial redundancy could be explained by the more acidic composition of the ATG2B FFNT motif compared to that of ATG2A, potentially enabling weak interactions with other VAP family members.

### MOSPD3 is required for ATG2A-mediated autophagosome biogenesis

Next, we investigated the functional significance of the ER recruitment of ATG2A by MOSPD3. First, we examined the role of ATG2A/B in autophagosome formation. We created ATG2A-KO, ATG2B-KO, and ATG2A/B-double-KO (DKO) HeLa cells using CRISPR–Cas9 (Fig. S4A-C). Immunoblotting showed that ATG2 DKO, but not ATG2A KO or ATG2B KO, caused a profound defect in autophagy, as shown by the strong accumulation of LC3B (Fig. S4D). This indicates that ATG2A and ATG2B act redundantly in autophagy, in line with the findings in previous studies^14, 28, 42^.

In view of such a robust defect in ATG2A/B-DKO cells, we sought to investigate the role of MOSPD3 in autophagosome biogenesis using a rescue strategy with the ATG2A/B-DKO cells. In the rescue experiments, cells positive for transfected GFP constructs were sorted by flow cytometry and subjected to immunoblotting (Fig. 5A). We found that the expression of GFP-ATG2B, but not GFP-ATG2A or its FFNT mutants T362A or T362D, significantly restored the autophagic defects, as indicated by the reduced LC3B level compared with that of ATG2 DKO (Fig. 5B). Two ATG2A mutants deficient in binding to IM (GFP-ATG2A-V1886E-I1890E-V1897E; GFP-ATG2A-F1758E-F1761E-L1765E)^28, 43^ did not rescue the defect, as was also the case for the coexpression of GFP-ATG2A and Halo-MOSPD3. In view of the LD localization of GFP-ATG2A, we concluded that ectopically expressed GFP-ATG2A may be dysfunctional in terms of autophagy.

**Fig. 5.**
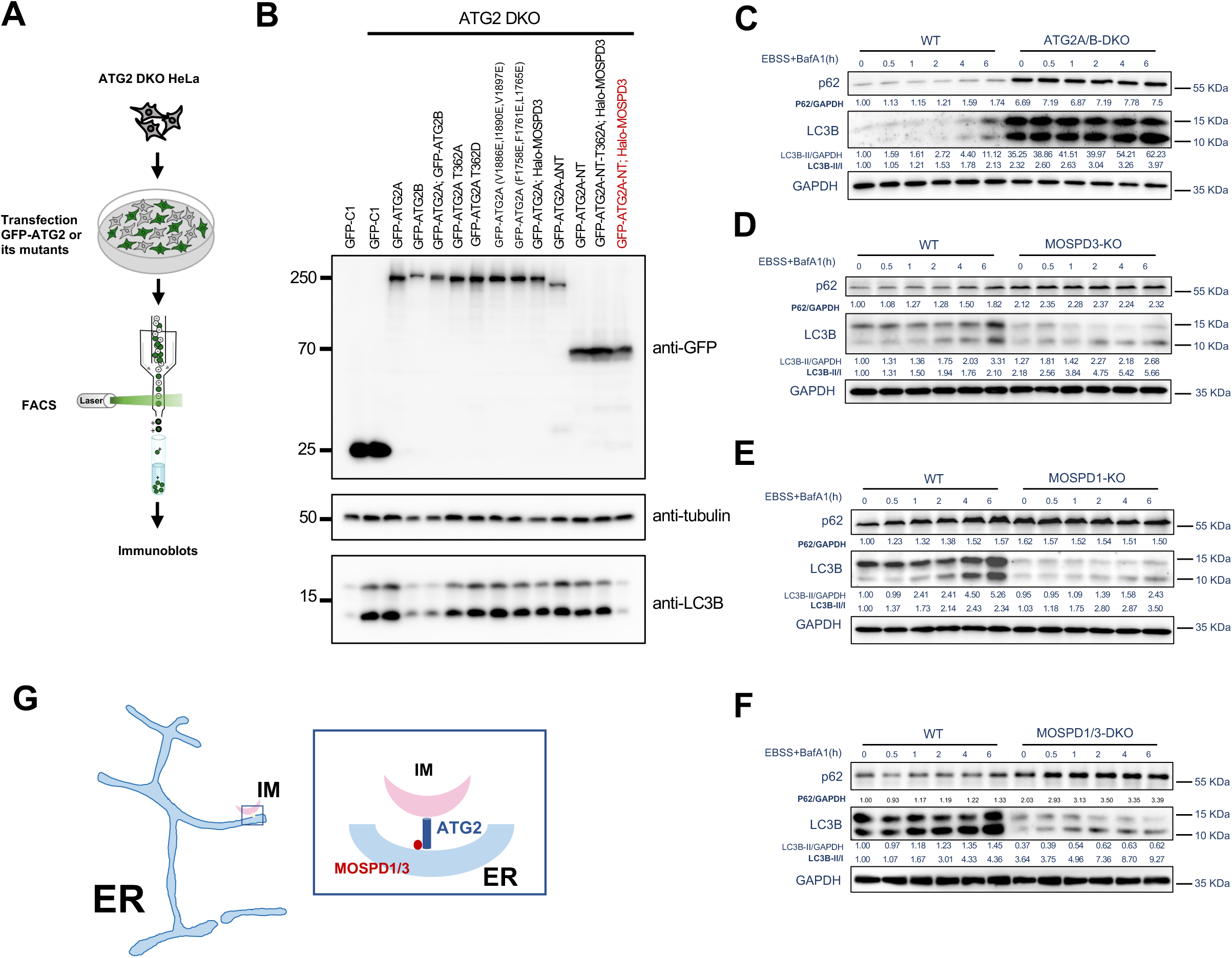
MOSPD3 is required for ATG2A-mediated autophagosome biogenesis. **a.** Workflow of Rescue experiment. GFP-ATG2A or its mutants were expressed in ATG2 DKO HeLa cells, followed by fluorescence-activated cell sorting (FACS) and immunoblot analysis. **b.** Western blots demonstrate the level of LC3B in rescue experiments as in (**a**) from 3 independent experiments. **c-f.** Western blots demonstrate the levels of p62 and LC3B in control, ATG2A/B-DKO (**c**), MOSPD3-KO (**d**), MOSPD1-KO (**e**) or MOSPD1/3 DKO (**f**) cells treated with EBSS (0, 0.5, 1, 2, 4, 6h) in the presence of BafA1 (1h) from 3 independent experiments. **g**. Working model for ATG2 and MOSPD1/3 at ER–IM contact sites. MOSPD3 acts as an ER adaptor for ATG2A, whereas multiple VAPs including MOSPD3 and MOSPD1 is responsible for ER recruitment of ATG2B in autophagy. The loss of MOSPD3 and/or MOSPD1 impedes autophagy.

Notably, GFP-ATG2A-NT, lacking the phagophore-targeting module at the C-terminal region, failed to rescue the defect, and an ATG2A truncation mutant lacking the NT region containing the FFNT motif (GFP-ATG2A-ΔNT) was also non-functional in the rescue experiment. However, coexpression of GFP-ATG2A-NT with Halo-MOSPD3, markedly rescued the autophagy defect (Fig. 5B). Importantly, coexpression of the FFNT motif mutant GFP-ATG2A-NT-T362A with Halo-MOSPD3 failed to rescue the defect, indicating an essential role of FFNT-MSP interaction in autophagy. Therefore, we conclude that the recruitment of ATG2A-NT to the ER via MOSPD3 is sufficient to restore the autophagic defect in ATG2A/B-DKO cells, suggesting that MOSPD3 plays a key role in ATG2A-mediated autophagosome formation.

In addition, we assessed autophagic flux in MOSPD3-deficient cells by examining the degradation of the selective autophagy adaptor p62/SQSTM1 upon starvation in presence of bafilomycin A1 (BafA1). As a control, ATG2 DKO resulted in strong accumulation of p62 and LC3 (Fig. 5C). Similarly, depletion of MOSPD3 (Fig. 5D), MOSPD1 (Fig. 5E) or both (MOSPD1/3 DKO; Fig. 5F) also causes accumulation of p62 compared to the control and this trend was more evident in the case of MOSPD1/3 DKO.

We found that, in these MOSPD protein-depleted cells, the LC3-II/LC3-I ratio was also increased compared to the control (Fig. 5C-E), similar to the phenotype induced by ATG2 DKO. Note that the LC3B mRNA levels were significantly decreased in these MOSPD KO cells (Fig. S4E), a finding that may reflect compensatory regulation and warrants further investigation.

Last, we investigated whether MOSPD1/3 deficiency has an effect on autophagosome formation. While depletion of either MOSPD3 or MOSPD1 exhibit a mild defect, double depletion of MOSPD3 and MOSPD1 caused a more profound defect in autophagosome formation, as revealed by higher colocalization between endogenous LC3B and the isolation membrane marker DFCP1 in MOSPD KO cells than in WT cells, similar to the dramatic colocalization of DFCP1 and LC3B in the ATG2 DKO cells (Fig. S4F, G), reflecting a block in autophagosome maturation. These results suggest a defect at a step downstream of IM initiation (e.g., progression of IMs into autophagosomes) in MOSPD1/3 KO, similar to the defect resulted from loss of ATG2A and ATG2B.

## Discussion

The source of membranes for autophagosome biogenesis has been debated for decades^2^. However, accumulating evidence now suggests that the ER is the main source of lipid supply for the growing autophagosomes, but the mechanism by which the core autophagy machinery ATG2 specifically recognizes the ER remains unclear, especially in mammals. In this study, we found that the atypical VAP protein MOSPD3 is an adaptor that recruits ATG2A to the ER. MOSPD3 colocalizes with ATG2A and is specifically enriched at ER sites adjacent to the IM during autophagosome formation. The recruitment depends on the interactions between the FFNT motif at the NT of ATG2A and an MSP domain of MOSPD3. Ectopic expression of MOSPD3 with ATG2A-NT, but not a MOSPD3-binding-defective mutant ATG2A-NT-T362A, markedly rescued the autophagic defect caused by ATG2A/B DKO. MOSPD3 depletion blocks the ER recruitment of ATG2A, and impairs autophagic flux (working model; Fig. 5G). Together, this study demonstrates that MOSPD3 functions as an ER adaptor for ATG2A in autophagosome biogenesis.

VAPA and VAPB (VAPA/B), two canonical VAP proteins, are known to play important roles in the early steps of autophagy, and VAPA/B interact with ULK1/FIP200 and WIPI2 and are involved in the formation and/or stabilization of the ER–IM contact that is essential for autophagosome formation^30^. Here, we identified that MOSPD1/3, two recently identified atypical VAP proteins, are also directly involved in autophagosome formation. In contrast to the role of VAPA/B in the stabilization of ULK1–FIP200 interaction and/or ER–IM contacts, our results indicate that MOSPD3 acts as an adaptor for the recruitment of ATG2A to the ER, a key step during IM membrane growth and autophagosome maturation. Interestingly, similar to the findings for MOSPD3 KO, VAPA/B depletion was reported to result in milder autophagy defects than deletion of ATG genes^30^, suggesting that both VAPA/B and MOSPD1/3 are involved in autophagy and that their functions may partially overlap, for example, in terms of the initiation and/or stabilization of ER–IM contacts as well as the ER recruitment of ATG2B. Therefore, in agreement with the previous study^30^, our results suggest diverse and crucial roles of VAPs in the early stage of autophagy.

FFNT motifs are FFAT-like motifs recently identified that have more neutral amino acids within them or nearby compared with the canonical FFAT sequence^44^. Notably, unlike the FFAT motif bound by VAPA/B or MOSPD2, FFNT motifs are recognized by MOSPD1/3^44^. Supporting this previous study, we found an FFNT motif in the NT of ATG2A and ATG2B that prefers MOSPD3 and MOSPD1 over other VAPs. Interestingly, ATG2A and ATG2B were indeed found among the protein candidates co-enriched with MOSPD1 and MOSPD3 by BioID in the previous study^44^. In addition, MOSPD1 and MOSPD3 can form a heterodimer^34^. However, our results showed that the mechanisms underlying the recruitment of ATG2A and ATG2B to the ER are different, with ATG2A recruitment being MOSPD3-dependent, whereas ATG2B recruitment is mediated by multiple VAPs including MOSPD3 and MOSPD1. A plausible explanation for this is that MOSPD1 and MOSPD3 may also form homodimers in cells, in addition to heterodimers, and that the MOSPD1 homodimer may be able to recruit ATG2B but not ATG2A to the ER. This hypothesis should be tested in future studies.

In contrast to the existence of two ATG2 paralogs in mammals, there is only one ATG2 gene in yeast, which encodes a single ATG2p protein. Interestingly, the FFNT motif of ATG2A or ATG2B is conserved in vertebrates but not in yeast, suggesting that ATG2 is recruited to the ER by other mechanisms in non-vertebrates. In support of this notion, yeast ATG2p was reported to bind to the ER exit site using basic residues in the NT and to the expanding edge of the IM through its C-terminal region, which was assisted by an Atg18–PtdIns3P interaction^13^.

It is noteworthy that VPS13 in yeast is recruited to multiple membrane contacts via different adaptors on different organelles^45^. In mammals, BLTP proteins including VPS13A^25, 46–51^, VPS13D^52–55^, SHIP164^56–58^ and BLTP2^59–61^ can be recruited to different types of membrane contacts under different conditions. A recent study showed that ATG2 can be recruited to endosome–IM contacts by a novel ATG2A-binding protein, ANKFY1, and mediates PI3P transfer from endosomes for IM expansion^29^. In addition, ATG2A has been shown to interact with the outer mitochondrial membrane protein TOM40 to promote IM expansion at ER–mitochondria contacts^62^. ATG2A is also reported to act as a tether during autophagosome-lysosome fusion, cooperated by WDR45 and WDR45B^63^. Besides, ATG2A is reported to be recruited to MCS between the ER and damaged lysosome for rapid lysosome repair^64^. Thus, ATG2 exhibits localization versatility, likely mediated by distinct adaptors under varying cellular conditions.

## Materials and methods

### Cell culture, transfection, RNAi

The African green monkey kidney fibroblast-like COS7 cell line (ATCC), human cervical cancer HeLa cells (ATCC), human embryonic kidney (HEK) 293T (ATCC) and human osteosarcoma U2OS ATG2A-KI cell line were grown in DMEM (Invitrogen) supplemented with 10% fetal bovine serum (Gibco). All of the cell lines used in this study were confirmed free of mycoplasma contamination.

Transfection of plasmids and RNAi oligos was carried out with Lipofectamine 2000 and RNAi MAX, respectively. For transfection, cells were seeded at 4 x 10^5^ cells per well in a six-well dish ∼16 h before transfection. Plasmid transfections were performed in OPTI- MEM (Invitrogen) with 2 μL Lipofectamine 2000 per well for 6 h, followed by trypsinization and replating onto glass-bottom confocal dishes at ∼3.5 x 10^5^ cells per well. Cells were imaged in live-cell medium (DMEM with 10% FBS and 20 mM Hepes with no penicillin or streptomycin) ∼16–24 h after transfection. For all transfection experiments in this study, the following amounts of DNA were used per 3.5 cm well (individually or combined for co-transfection): 100 ng for GFP-MOSPD1/3, Halo-MOSPD1/3 and its mutants, 500 ng for GFP-ATG2A/B and its mutants; 500 ng for other VAP proteins and TMEM41B and VMP1; 50 ng for GFP-DFCP1; 500 ng for mCh-LC3B; 500 ng for Halo-Sec61β. For siRNA transfections, cells were plated on 3.5 cm dishes at 30–40% density, and 2 μl Lipofectamine RNAimax (Invitrogen) and 50 ng siRNA were used per well. At 48 h after transfection, a second round of transfection was performed with 50 ng siRNAs. Cells were analysed 24 h after the second transfection for suppression. All siRNAs used in all experiments were synthesized by RiboBio. siRNA targeting sequences for human VMP1#1: GGCACAAAGTTATGCCAAA; human VMP1#2: GTGCAACCCTAATTGGAAA; human VMP1#3: GGAGTATCATCTCAAAAGT; human TMEM41B#1: GCTGGATCAGCAAGAATGT; human TMEM41B #2: TCAAGTACTTGTAGCTTAT; human TMEM41B #3: CCAAGGCTCTAGGAAAAGT.

### Plasmids

ATG2A (NM_015104), ATG2B (NM_018036), MOSPD3 (NM_001040097), MOSPD1 (NM_001100814.3), MOSPD2, VAPA, VAPB, TMEM41B, VMP1, and DFCP1 were cloned from HeLa cDNA. The ORFs of these genes were cloned into mEGFP-C1/Halo-C1 vector. All mutant constructs in this study were generated by PCR mediated site directed mutagenesis on the basis of their respective wild-type constructs. All constructs were confirmed by DNA sequencing. A list of plasmids and primers for DNA constructs were provided in Table 1 and 2, respectively.

**Table 1.** Plasmids used in this study.

| Plasmid | Source |
| --- | --- |
| mEGFP-C1 | addgene 54579 |
| mEGFP-N1 | addgene 54767 |
| Halo-LC3B | This study |
| GFP-ATG2A | This study |
| Halo-VAPA | This study |
| SNAP-VAPB | This study |
| Halo-MOSPD1 | This study |
| Halo-MOSPD2 | This study |
| Halo-MOSPD3 | This study |
| GFP-ATG2A-N | This study |
| Halo-VMP1 | This study |
| TMEM41B-Halo | This study |
| Halo-ATG2A-ΔFFNT | This study |
| GFP-ATG2A T362A | This study |
| GFP-ATG2A T362D | This study |
| GFP-ATG2B | This study |
| GFP-ATG2B-ΔFFNT | This study |
| Halo-MOSPD3-ΔMSP | This study |
| Halo-MOSPD3-ΔTM | This study |
| Halo-MOSPD3-ΔPro | This study |
| Halo-MOSPD3-MSP | This study |
| mCherry-LC3B | This study |
| GFP-DFCP1 | This study |
| GFP-MOSPD3 | This study |
| 14xHis-ATG2A | This study |
| 14xHis-ATG2B | This study |
| GFP-ATG2A (F1758E, F1761E, L1765E) | This study |
| GFP-ATG2A (V1886E, I1890E, V1897E) | This study |

**Table 2.**
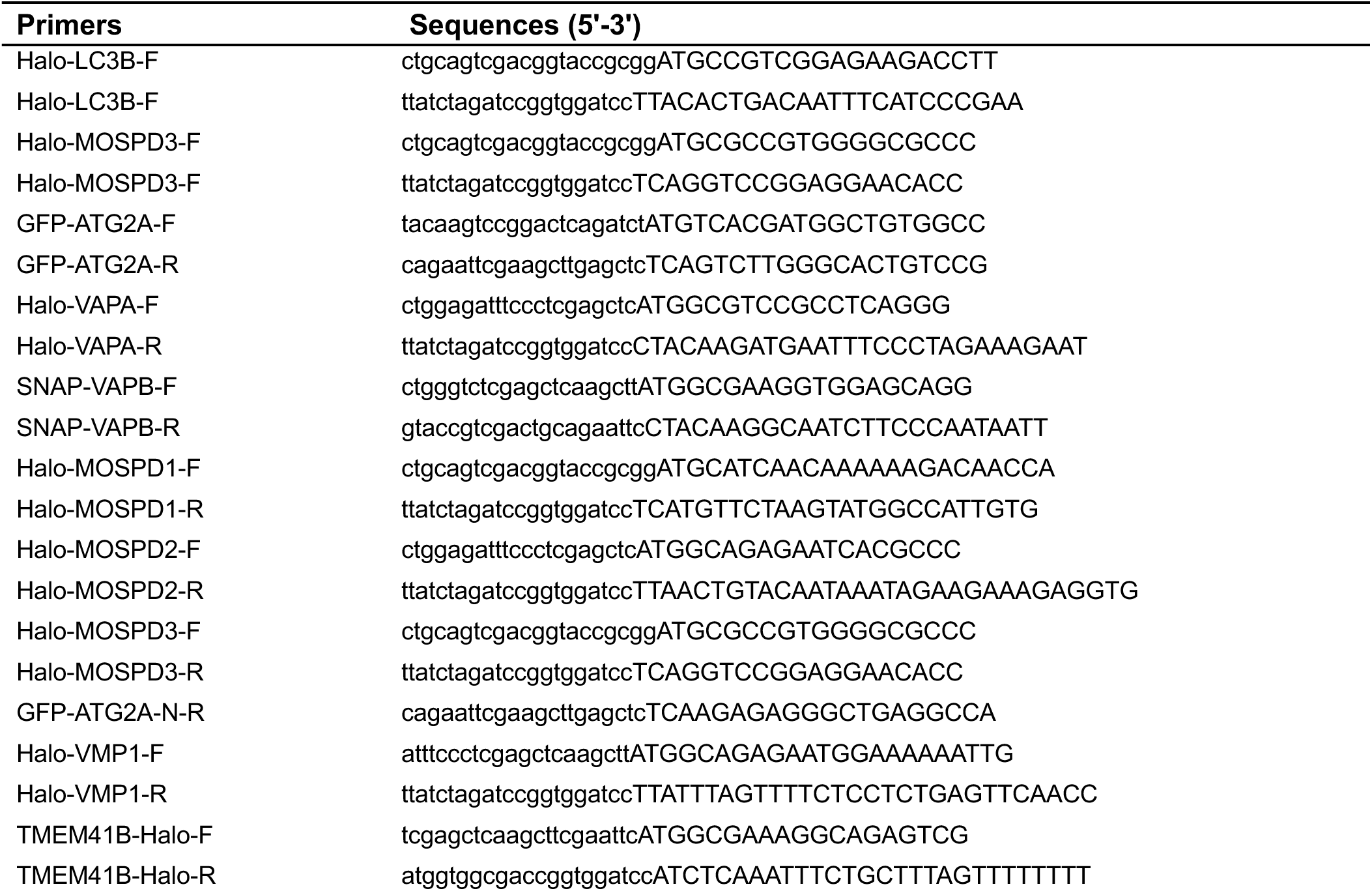
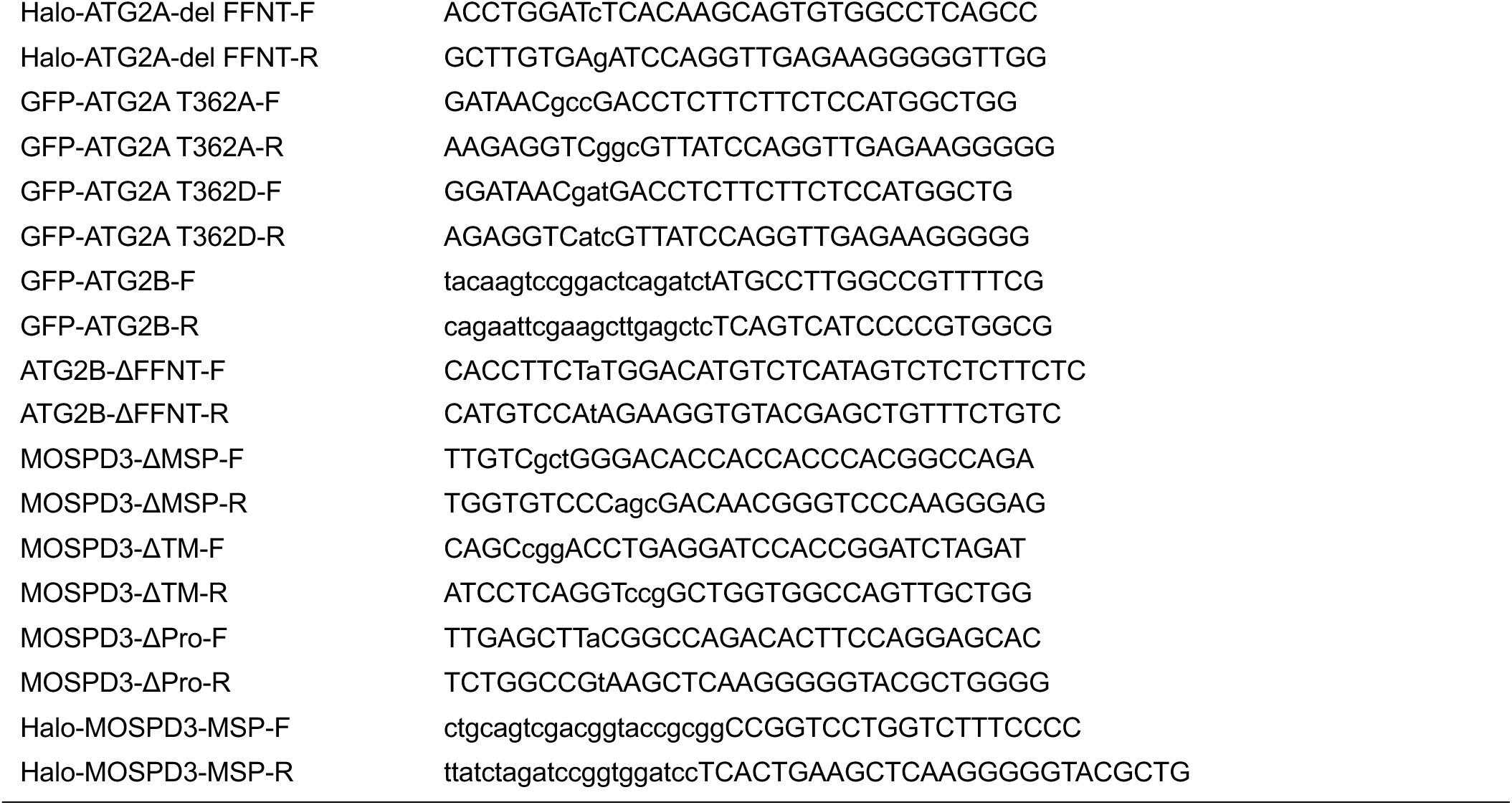
Primers used in this study.

### Antibodies and reagents

Anti-ATG2A (15011, CST), anti-ATG2B (A8498, Abclonal) and anti-MOSPD3 (C-6) (sc-514923, Santa cruz) were used at 1:1000 for western blots. Anti-Flag (F1804, Sigma-Aldrich) were used at 1:1500 for western blots. Anti-Calnexin (ab22595, Abcam), anti-PMP70 (ab3421, Abcam), anti-LAMP1(H4A3) (sc-20011, Santa cruz), anti-TOM20 (sc17764, Santa cruz), anti-LC3B (A19665, ABclonal) and anti-GM130 mAb (M179-3, MBL) were used at 1:2000 for western blots. Anti GFP-Tag pAb (AE011, ABclonal), anti-Halo Tag pAB (G928A, Promega), anti-GAPDH (AB9485, Abcam), anti-tubulin (100109-MM05T, Sino Biological Inc), anti GST-Tag mAb (AE001, Abclonal) and anti-His (66005-1-Ig, Proteintech) were used at 1:5000 for western blots. Anti-phospho-threonine an7body (AB1607, Sigma) were used at 1:200 for western blots. For immunoblotting, HRP-conjugated goat anti-mouse IgG (H+L) (AS003, Abclonal), HRP-conjugated goat anti-rabbit IgG (H+L) (AS014, Abclonal) were used as secondary antibodies.

For live cell imaging, Janilia Fluo^®^ 646 HaloTag^®^ Ligand (GA1120, Promega) and BODIPY-558/568 C12 (D3835, ThermoFisher Scientific) were added to OPTI-MEM (Invitrogen) medium in concentration of 20 nM, followed by incubation for 20 min. Cells were washed three times with PBS before imaging.

### CRISPR-Cas9-mediated gene editing

To make ATG2A KO HeLa cell lines, two gRNAs (5^’^-tctttcacacagtttgacca-3^’^ and 5^’^-cctggaaatctgggtgagga-3^’^) were used to delete 137 bp from exon 1 of ATG2A gene. To make ATG2B KO HeLa cell lines, two gRNAs (5^’^-GCGGGGCCTAAGCCTGGGG-3^’^ and 5^’^-AAATGGGTAAGAGCTGCCGG-3^’^) were used to delete exon 1 of ATG2B gene. To make MOSPD1 KO HeLa cell lines, two gRNAs (5^’^-tgcataaaatatgagctccg-3^’^ and 5^’^-aagtttccgagcaaagccaa-3^’^) were used to delete exon 1-3 of MOSPD1 gene. To make MOSPD3 KO HeLa cell lines, two gRNAs (5^’^-cccggatctagtattcaggg^’^ and 5^’^-cttcctcagacagctcaatg-3^’^) were used to delete exon 1-3 of MOSPD3 gene. Complementary gRNAs were annealed and subcloned into the pSpCas9(BB)-2A-GFP (pX-458) vector (#48138; Addgene) between BbsI endonuclease restriction sites. Upon transfection, HeLa cells were grown in an antibiotic-free medium for 48 h, followed by single-cell sorting by fluorescence-based flow cytometry. Two independent clones were verified by imaging and western blots.

### Live imaging by high-resolution confocal microscopy

Cells were grown on 35 mm glass-bottom confocal MatTek dishes, and the dishes were loaded to a laser scanning confocal microscope (LSM900, Zeiss, Germany) equipped with multiple excitation lasers (405 nm, 458 nm, 488 nm, 514 nm, 561 nm and 633 nm) and a spectral fluorescence GaAsP array detector. Cells were imaged with the 63×1.4 NA iPlan-Apochromat 63 x oil objective using the 405 nm laser for BFP, 488 nm for GFP, 561nm for OFP, tagRFP or mCherry and 633nm for Janilia Fluo® 646 HaloTag® Ligand.

### GFP-trap assay

GFP trap (GTA-100, ChromoTek) was used for detection of protein–protein interactions and the GFP-Trap assays were performed according to the manufacturer’s protocol. 5% input was used in GFP traps unless otherwise indicated.

### GST Tag and His Tag protein purification

GST and His constructs were transformed into Escherichia coli BL21 (DE3) cells, and cells were incubated at 37°C until the optical density (OD) at 600 nm reached 0.6–0.8. Subsequently, cells were incubated at 16°C for another hour, followed by induction with 1 mM IPTG overnight at 16°C. Cells were lysed via sonication. GST fusion proteins were purified via the GST-tag Protein Purification kit (C600031-0025, Sangon, China), His fusion proteins were purified via the Ni-NTA Sefinose (TM) Resin Purification kit (G600033-0100, Sangon, China).

### In vitro Pull-down assays

HEK293 cells transiently transfected with GFP-MOSPD3 were lysed in high-salt lysis buffer (RIPA buffer containing 500 mM NaCl, proteasome inhibitors and PMSF). GFP-Trap beads were used to pellet GFP-MOSPD3 or GFP tag only from cell lysates, followed by washing with high-salt lysis buffer for 10 times. The GFP beads were incubated with purified 14xHis–ATG2A-NT or 14xHis–ATG2B-NT overnight at 4°C, respectively, followed by washing beads with freshly prepared HNM buffer (20 mM Hepes, pH 7.4, 0.1 M NaCl, 5 mM MgCl2, 1 mM DTT and 0.2% NP-40). The GFP beads were resuspended in 100 μL 2 x SDS-sampling buffer. Re-suspended beads were boiled for 10 min at 95°C to dissociate protein complexes from beads. Western blotting was performed using anti-GFP or His antibodies. The Coomassie staining was performed for purified 14xHis–ATG2A-NT or 14xHis–ATG2B-NT.

### Purification of ER membranes by density gradient centrifugation

ER fractions were enriched using Endoplasmic Reticulum Isolation Kit (Sigma-Aldrich, ER0100) according to the manufacturer’s instructions. Briefly, HeLa cells from five confluent 10-cm dishes were collected, followed by centrifugation at 600 x g for 5 min. After washing the cells three times with PBS, the packed cell volume (PCV) was measured and then suspended in a volume of Hypotonic Extraction Buffer (10 mM HEPES, pH 7.8, with 1 mM EGTA and 25 mM potassium chloride) equivalent to 3 times the PCV. After the incubation of the cells for 20 minutes at 4 °C allowing the cells to swell, the cells were centrifuged at 600xg for 5 minutes, followed by the measurement of the “new” PCV. After adding a volume of Isotonic Extraction Buffer (10 mM HEPES, pH 7.8, with 0.25 M sucrose, 1 mM EGTA, and 25 mM potassium chloride) equivalent to 2 times the “new” PCV, the suspension was then transferred to a 7-mL Dounce homogenizer, followed by the lysis of the cells with 10 strokes and then the centrifugation of the homogenate at 1,000 x g for 10 min at 4 °C. After the transfer of the supernatant to another centrifuge tube, the supernatant was centrifuged at 12,000 x g for 15 min at 4 °C, followed by another centrifugation for 60 min at 100,000 x g at 4 °C. The pellet was the microsomal fraction and further verified by western blots using anti-Calnexin antibody.

### Statistical analysis

All statistical analyses and p-value determinations were performed in GraphPad Prism6. All the error bars represent Mean ± SD. To determine p-values, ordinary one-way ANOVA with Tukey’s multiple comparisons test was performed among multiple groups and a two-tailed unpaired student t-test was performed between two groups.

## Acknowledgements

We thank Qing Zhong (Shanghai Jiao Tong University) for sharing the U2OS ATG2A-KI cell line. We thank Tom Buckle from Scribendi (www.scribendi.com) for editing a draft of this manuscript. We thank the School of Basic Medicine Innovation Research Center, Huazhong University of Science and Technology, for providing technical supports. W. Ji was supported by National Natural Science Foundation of China (32570822, 32371343, 32541022).

The authors declare no competing interests.

## Author contributions

Y. Du and W. Ji conceived the project and designed the experiments. Y. Du, T. Zhou, Y. Liu, N. Ye, B. Zou, W. Chang, and L. Deng performed the experiments. Y. Du, T. Zhou, Y. Liu, A. Shi, and W. Ji analysed and interpreted the data. W. Ji prepared the manuscript with inputs and approval from all authors.

## Data and materials availability

All the data and relevant materials, including reagents and primers, that supports the findings of this study are available from the corresponding author upon reasonable request.

---

**Supplementary Fig. 1.**
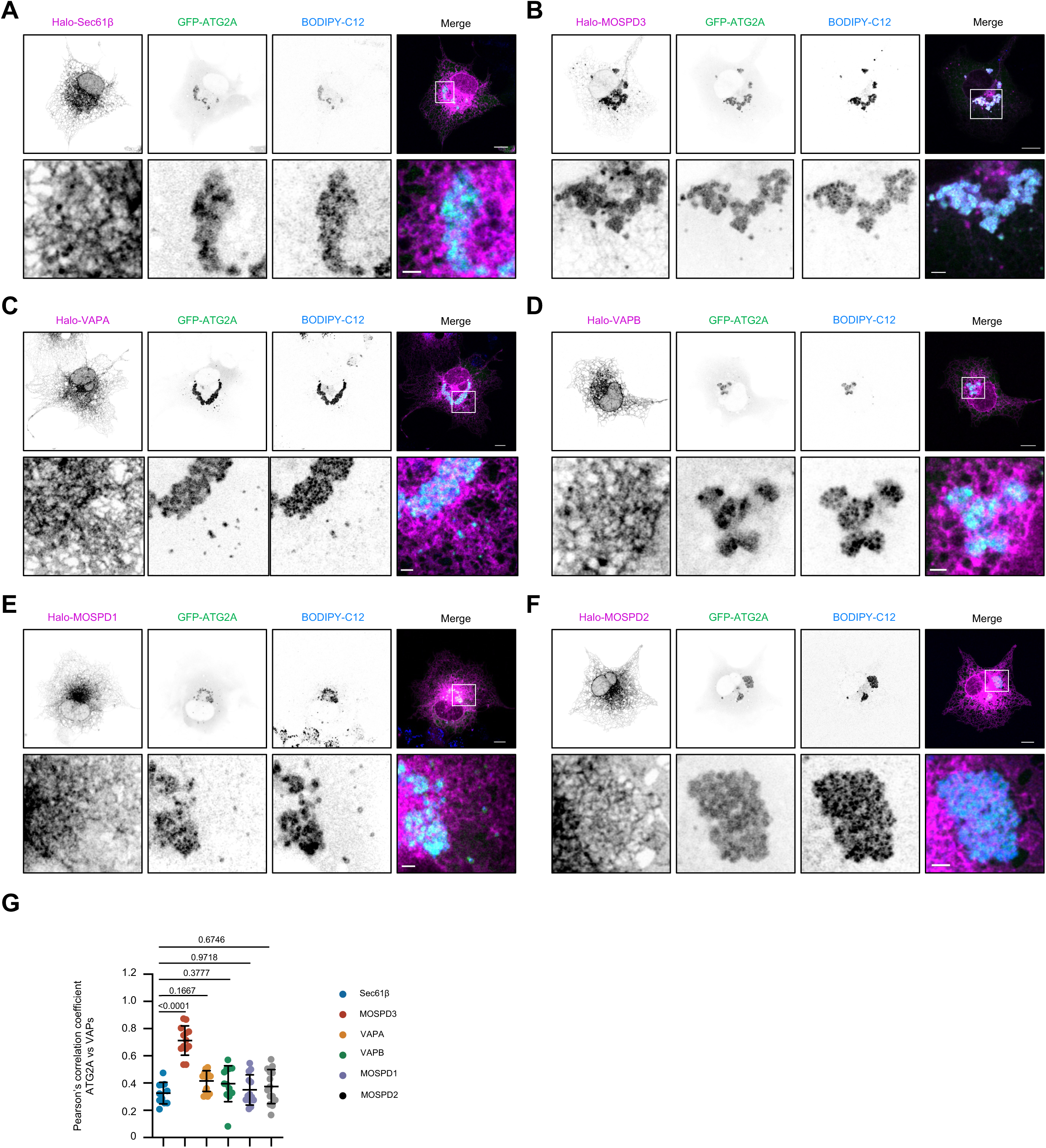
The colocalization between GFP-ATG2A and Halo-VAPs. **a-f**. Representative images of live BODIPY (blue)-stained COS7 cells transiently transfected with GFP-ATG2A (green) and Halo-VAPs (magenta; MOSPD1/2/3, VAPA and VAPB) with insets from at least 3 independent experiments. **g.** In cells as in (**a-f**), Pearson’s correlation coefficient of ATG2A vs different ER proteins including Halo-Sec61β (11 cells), Halo-MOSPD1 (13 cells), Halo-MOSPD3 (13 cells), Halo-MOSPD2 (14 cells), Halo-VAPA (13 cells), or Halo-VAPB (12 cells) in three independent experiments. Ordinary one-way ANOVA with Tukey’s multiple comparisons test. Mean ± SD. Scale bar, 10 μm in the whole cell images and 2 μm in the insets in (a-f).

**Supplementary Fig. 2.**
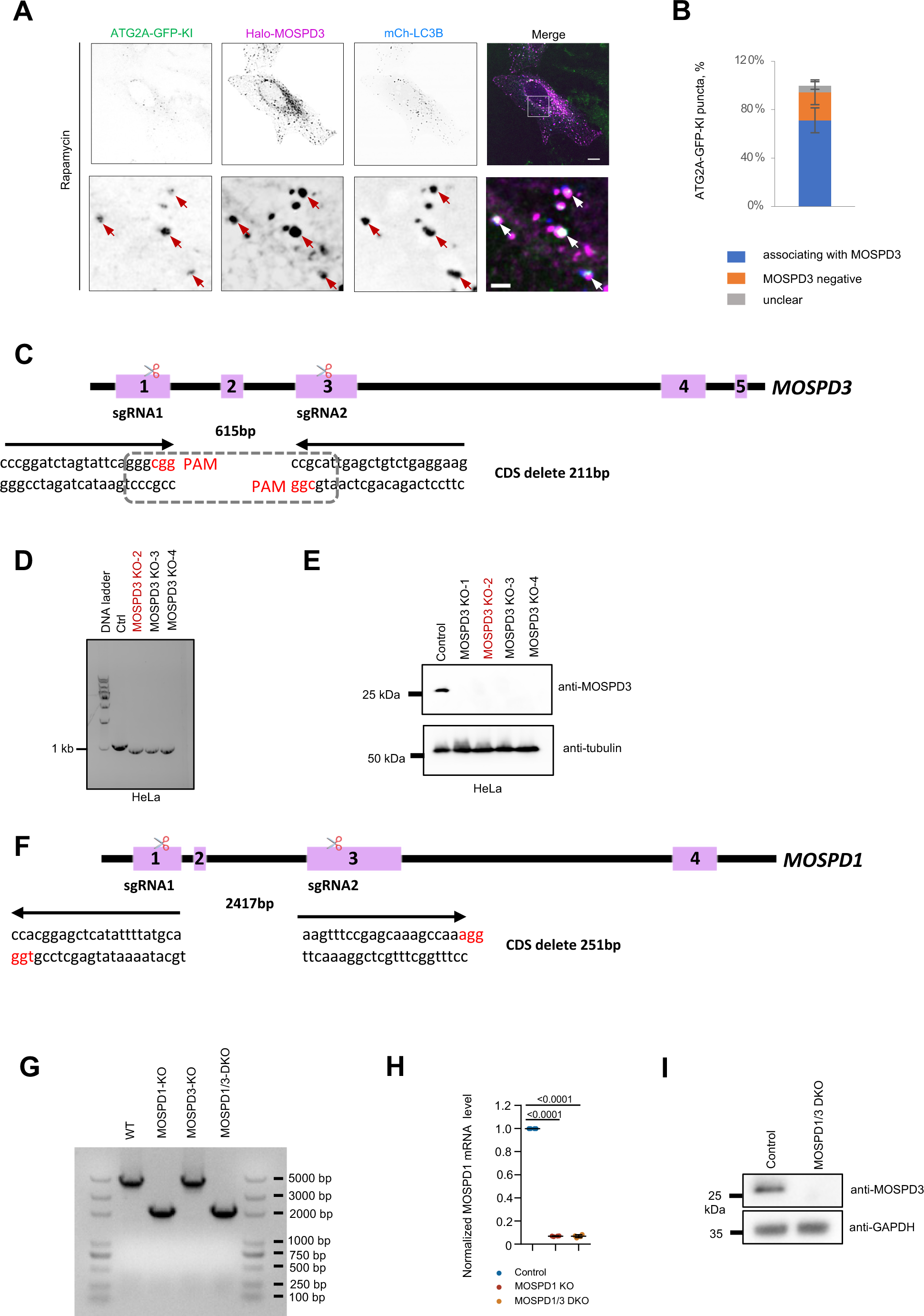
MOSPD3 and MOSPD1 KO HeLa cells. **a.** Representative images of a live U2OS ATG2A-GFP-KI cell transiently transfected with Halo-MOSPD3 (magenta) and mCh-LC3B (blue) with insets under Rapamycin stimulation. **b.** The distribution of endogenous ATG2A puncta relative to MOSPD3 puncta (540 ATG2A-GFP puncta from 12 cells) from at least 3 independent experiments. Mean ± SD. **c.** CRISPR knock-out of MOSPD3 in HeLa cells (MOSPD3-KO). Two sgRNAs are used with the lines above the sequence, indicating the PAMs (CGG for sgRNA1 and CGG for sgRNA2) for spCas9. **d.** Genotyping of control and three MOSPD3 KO clones (#2, #3, and #4) by DNA gel electrophoresis. **e.** The depletion of MOSPD3 in four MOSPD3-KO clones (#1, #2, #3 and #4) were confirmed by immunoblots. The KO clone #2 was used in the study. **f.** CRISPR knock-out of MOSPD1 in HeLa cells (MOSPD1-KO). Two sgRNAs are used with the lines above the sequence, indicating the PAMs (TGG for sgRNA1 and AGG for sgRNA2) for spCas9. **g.** Genotyping of the MOSPD1 gene locus from control, MOSPD1 KO, MOSPD3 KO and MOSPD1/3 DKO by DNA gel electrophoresis using a pair of primers as indicated in (**f**). **h.** The mRNA levels of MOSPD1 in control, MOSPD1-KO, or MOSPD1/3-DKO HeLa cells were measured by qPCR based on three repeats. Mean ± SD. **i.** The depletion of MOSPD3 in the MOSPD1/3-DKO clones were confirmed by immunoblots. Scale bar, 10 μm in the whole cell images and 2 μm in the insets in (a).

**Supplementary Fig. 3.**
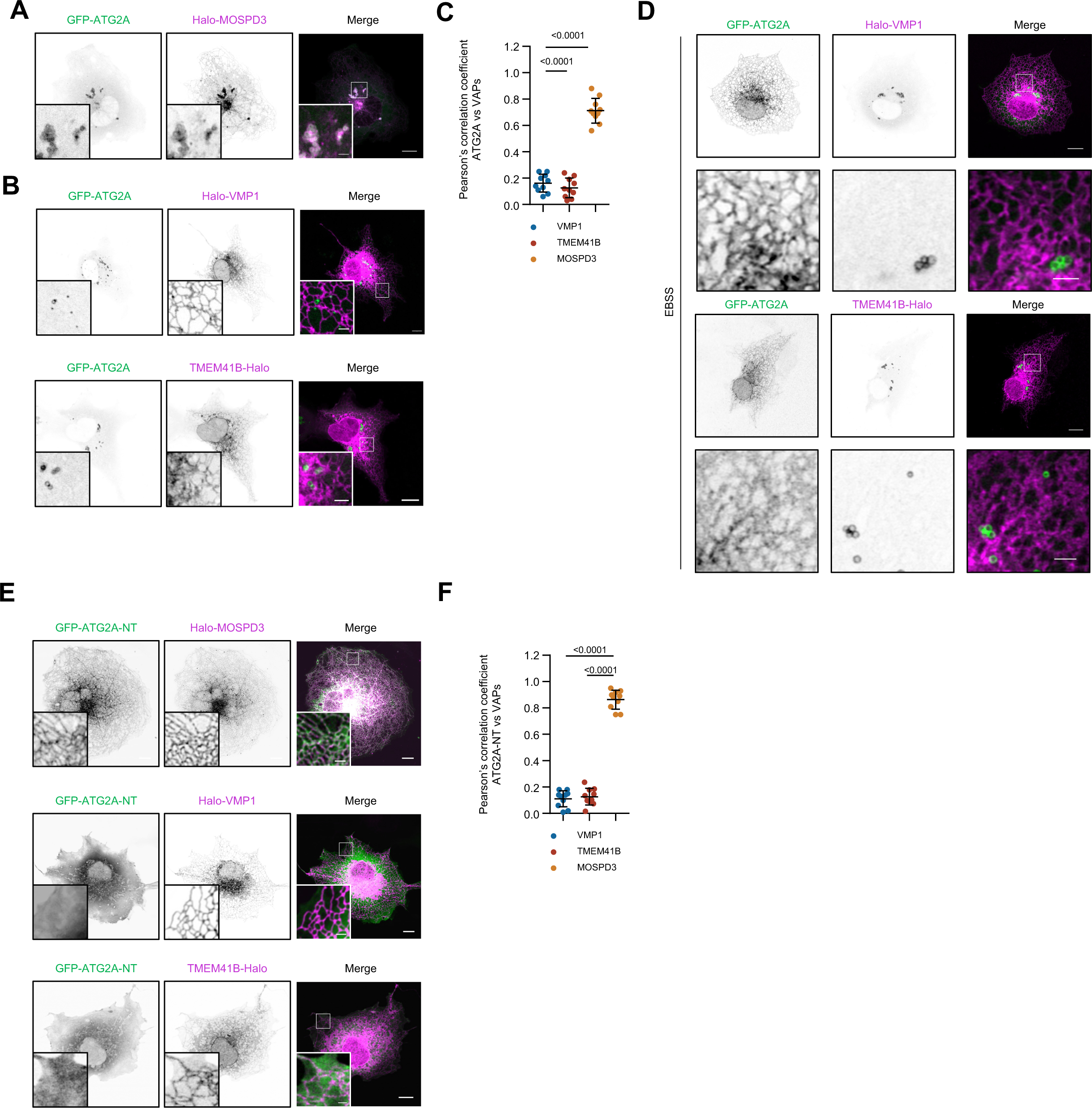
The colocalization between ATG2A and VPM1 or TMEM41B. **a, b**. Representative images of COS7 cells expressing GFP-ATG2A (green) along with either Halo-MOSPD3 (magenta; **a**) or Halo-VMP1 or TMEM41B (magenta; **b**) with insets. **c.** In cells as in (**a, b**), Pearson’s correlation coefficient of ATG2A vs TMEM41B (10 cells), VMP1 (10 cells) or MOSPD3 (10 cells) in three independent experiments. Ordinary one-way ANOVA with Tukey’s multiple comparisons test. Mean ± SD. **d.** Representative images of COS7 cells expressing GFP-ATG2A (green) along with either Halo-VMP1 (magenta; top panel) or TMEM41B (magenta; bottom panel) with insets upon EBSS starvation from three independent experiments. **e.** Representative images of COS7 cells expressing GFP-ATG2A-NT (green) with Halo-MOSPD3, Halo-VMP1, or TMEM41B-Halo (magenta) with insets. **f.** In cells as in (**e**), Pearson’s correlation coefficient of ATG2A-NT proteins vs VMP1 (10 cells), TMEM41B (10 cells), or MOSPD3 (10 cells) from three independent experiments. Ordinary one-way ANOVA with Tukey’s multiple comparisons test. Mean ± SD. Scale bar, 10 μm in the whole cell images and 2 μm in the insets in (a, b, d & e).

**Supplementary Fig. 4.**
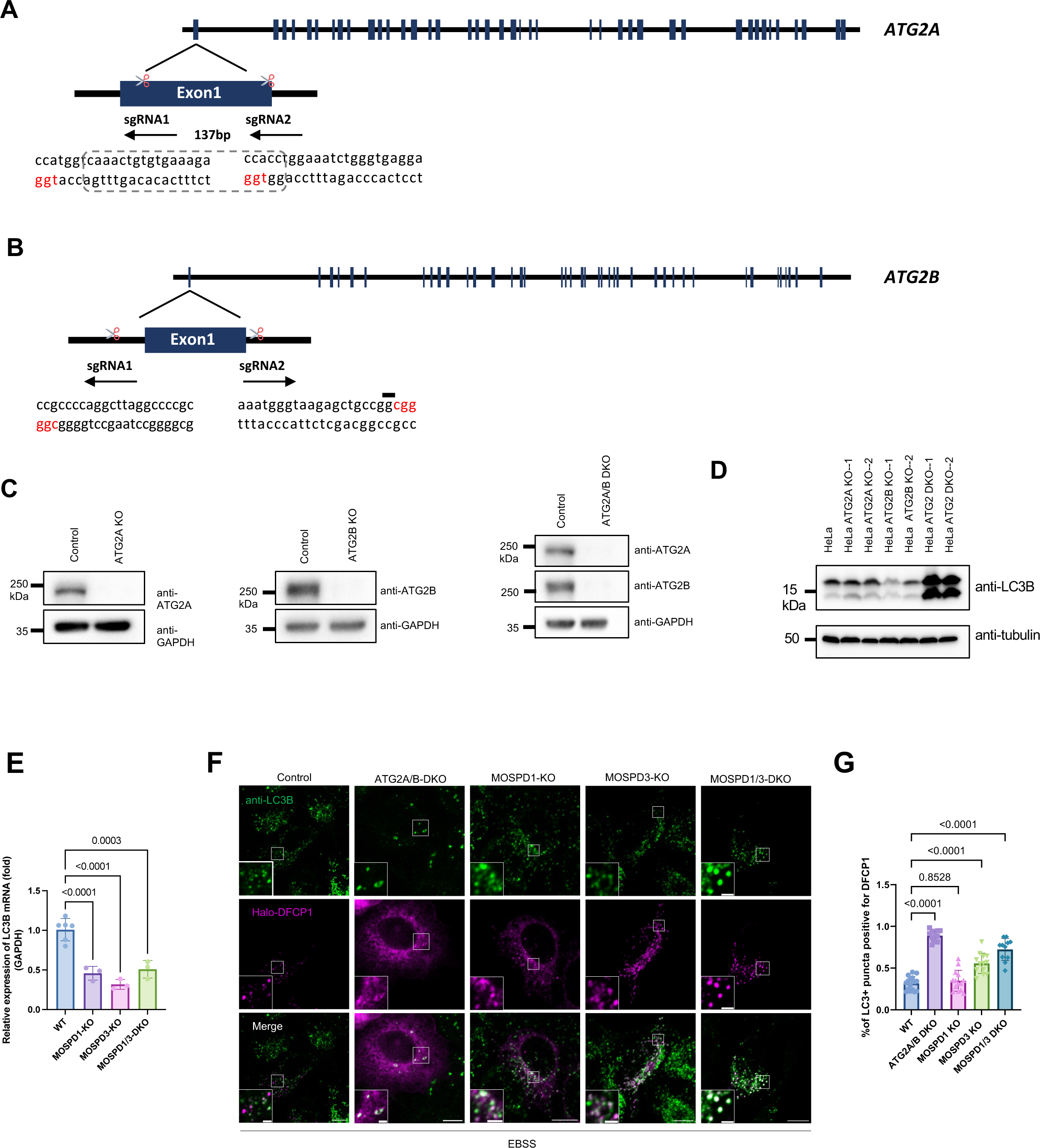
ATG2A/B DKO HeLa cell line and a role of MOSPD 1/3 proteins in autophagy. **a.** CRISPR knock-out of ATG2A in HeLa cells (ATG2A-KO). Two sgRNAs are used with the lines above the sequence, indicating the PAMs (TGG for sgRNA1 and TGG for sgRNA2) for spCas9. **b.** CRISPR knock-out of ATG2B in HeLa cells (ATG2B-KO). Two sgRNAs are used with the lines above the sequence, indicating the PAMs (CGG for sgRNA1 and CGG for sgRNA2) for spCas9. **c.** Western blots confirming the depletion of ATG2A or ATG2B in ATG2A KO (left panel), ATG2B KO cell line (middle panel), or ATG2A/B DKO cells (right panel). **d.** Western blots showing LC3B levels in control, ATG2A KO, ATG2B KO, and ATG2A/B-DKO cells. **e.** The mRNA levels of LC3B in control, MOSPD1-KO, or MOSPD1/3-DKO HeLa cells were measured by qPCR based on three repeats. Mean ± SD. **f.** Representative images of control, ATG2A/B DKO, MOSPD1 KO, MOSPD3 KO, MOSPD1/3 DKO HeLa cells stained with antibodies against LC3B (green) and expressing Halo-DFCP (magenta) with insets. **g.** Quantification of LC3B puncta relative to DFCP1 in cells as shown in (**f**). More than 12 cells were analysed from three independent experiments. Ordinary one-way ANOVA with Tukey’s multiple comparisons test. Mean ± SD.

## Notes

### Competing Interest Statement

The authors have declared no competing interest.

